# Soft sweeps are the dominant mode of adaptation in the human genome

**DOI:** 10.1101/090084

**Authors:** Daniel R. Schrider, Andrew D. Kern

**Affiliations:** Department of Genetics, Rutgers University, Piscataway, NJ, 08854, USA; Human Genetics Institute of New Jersey, Rutgers University, Piscataway, NJ, 08554, USA

**Keywords:** adaptation, selective sweeps, soft sweeps, machine learning, population genomics

## Abstract

The degree to which adaptation in recent human evolution shapes genetic variation remains controversial. This is in part due to the limited evidence in humans for classic “hard selective sweeps,” wherein a novel beneficial mutation rapidly sweeps through a population to fixation. However, positive selection may often proceed via “soft sweeps” acting on mutations already present within a population. Here we examine recent positive selection across six human populations using a powerful machine learning approach that is sensitive to both hard and soft sweeps. We found evidence that soft sweeps are widespread and account for the vast majority of recent human adaptation. Surprisingly, our results also suggest that linked positive selection affects patterns of variation across much of the genome, and may increase the frequencies of deleterious mutations. Our results also reveal insights into the role of sexual selection, cancer risk, and central nervous system development in recent human evolution.

## INTRODUCTION

Spurred by the ongoing revolution in DNA sequencing capacity, human population genetic datasets have grown exponentially in size over the past five years (Auton et al. 2015; UK10K Consortium 2015). Such growth enables insight into the evolutionary histories of human populations with hitherto unrivaled precision. A central question in the study of human evolution is the extent to which adaptation has driven recent evolution and affected patterns of genetic diversity (Akey 2009). This can be addressed by scanning genomic data for evidence of selective sweeps, wherein a beneficial mutation is favored by natural selection and therefore rapidly increases in frequency within a population. Such selective sweeps leave a characteristic footprint in variation; they create a valley of diversity around the selected site (Maynard Smith and Haigh 1974; Kaplan et al. 1989; Stephan et al. 1992), a deficit of both low- and high-frequency derived alleles at linked sites (Fay and Wu 2000), and an increase in linkage disequilibrium in flanking regions (Kim and Nielsen 2004). Thus there are multiple population genetic signals to exploit. Accordingly numerous theoretical and methodological advances (Kaplan et al. 1989; Stephan et al. 1992; Fu 1997; Kim and Stephan 2002; Nielsen et al. 2005b; Voight et al. 2006) in the study of selective sweeps have given researchers the ability to uncover the genetic basis of adaptation on a genome-wide scale.

There are two complimentary approaches to studying the impact of adaptive evolution on genetic variation. The first approach aims to infer genome-wide rates of adaptive evolution by estimating the mean effects of selective sweeps across the genome (Wiehe and Stephan 1993; Kern et al. 2002; Andolfatto 2007; Jensen et al. 2008; Hernandez et al. 2011; Sattath et al. 2011). Such approaches may estimate the rates of sweeps or their effects with respect to the genomic background, but do not focus on the targets of sweeps themselves. An alternative approach is to focus on finding individual selective sweeps throughout the genome, and in so doing characterize specific cases of adaptation with hopes of gaining general insight into the adaptive process (Sabeti et al. 2002; Voight et al. 2006; Williamson et al. 2007). The search for selective sweeps has shed light into the recent evolutionary histories of natural populations, and has shown a pervasive impact of adaptive evolution on polymorphism in some species such as *Drosophila melanogaster* (Begun et al. 2007; Macpherson et al. 2007; Langley et al. 2012; Lee et al. 2013; Garud et al. 2015). In humans, the picture remains less clear: while scans for selective sweeps have discovered numerous compelling candidates for strong positive selection (e.g. Ruwende et al. 1995; Stephens et al. 1998; Tishkoff et al. 2007; Bryk et al. 2008; Huerta-Sánchez et al. 2014), some recent studies have suggested that the impact of adaptation on patterns of variation genome-wide is quite limited (Hernandez et al. 2011; Lohmueller et al. 2011). Conversely, Enard et al. (2014) argue that the genome-wide reduction in diversity around substitutions is driven in part by positive selection.

One possible explanation for the difficulty in characterizing the contributions of adaptive and non-adaptive forces in human populations is that genetic hitchhiking effects may be muted by human demographic history. Many human populations appear to have experienced bottlenecks and/or recent growth (Marth et al. 2004; Fagundes et al. 2007; Gravel et al. 2011; Auton et al. 2015), which cause much of the genome to resemble selective sweeps (Nielsen et al. 2005b). Moreover, positive selection has historically been modeled as the process of a *de novo* beneficial mutation rapidly sweeping to fixation, a process now referred to as a hard sweep. However selection may act on previously segregating neutral or weakly deleterious variants (Orr and Betancourt 2001; Innan and Kim 2004). Selection on standing variation will produce qualitatively different skews in linkage disequilibrium and allele frequencies, along with a shallower valley in diversity (Hermisson and Pennings 2005; Przeworski et al. 2005; Berg and Coop 2015; Schrider et al. 2015)—such an event is thus referred to as a soft sweep. If selection typically proceeds through soft sweeps, as may be the case in Drosophila (Garud et al. 2015), then many sweeps may have been missed by previous scans that were designed to detect signatures produced under a hard sweep model.

We sought to address the controversy over the impact of adaptation on human genomic variation by conducting a genome-wide scan for both hard and soft selective sweeps across human populations. We previously developed S/HIC (Soft/Hard Inference through Classification), a machine learning method capable of detecting completed sweeps and inferring their mode of selection with unparalleled accuracy and robustness to non-equilibrium demography (Schrider and Kern 2016). Here we apply S/HIC to uncover hard and soft sweeps in six population samples from the 1000 Genomes Project (Auton et al. 2015), thereby performing the most comprehensive investigation of completed selective sweeps in humans to date. Surprisingly, our results suggest that patterns of polymorphism across much of the human genome may be affected by linked positive selection—primarily soft sweeps. Moreover, we find evidence that the mode of selection differs substantially across populations, with non-African populations adapting via hard sweeps to a much greater extent than African populations. Finally, we investigate the biological targets of selection in recent human evolution, with particular processes such as immunity, cancer, and sexual reproduction playing outsized roles.

## RESULTS

We set out to detect completed hard and soft selective sweeps in six populations from Phase 3 of the 1000 Genomes Project: two West-African populations (YRI and GWD from Yoruba and The Gambia, respectively), one East-African population (LWK from Kenya), one European population (CEU, from Utah, USA), one East Asian population (JPT from Japan), and one from the Americas (PEL from Peru). For each population we trained and applied a S/HIC classifier to identify hard and soft selective sweeps across the genome (Methods), distinguishing them from neutrally evolving regions as well as those linked to sweeps (Schrider and Kern 2016). Briefly, S/HIC is a machine learning method that leverages spatial patterns of a variety of statistics across a large genomic window in order to infer the mode of evolution at the center of the window. We previously showed that S/HIC is exceptionally robust to the confounding effect of linked selection (e.g. the “soft shoulder” effect where regions linked to hard sweeps resemble soft sweeps; Schrider et al. 2015), as well as non-equilibrium demographic histories, making it well suited for a survey of positive selection in humans. We also assessed the accuracy of our classifiers on simulated test data with the same demographic history used to generate training data, finding that S/HIC achieved good power for each demographic history, with somewhat higher accuracy for histories inferred from the African than non-African populations (supplementary fig. S1).

We also performed forward simulations under the GWD and JPT models (Methods) in order to assess whether purifying selection and its effect on variation at linked unselected sites (i.e. background selection Charlesworth et al. 1993) could result in false sweep calls. The results of these simulations suggest that S/HIC’s false positive rate is essentially unaffected by these forces (supplementary fig. S1). Note that we exposed each classifier to a wide range of mutation and recombination rates (see Methods) during training (and testing) in order to improve (and assess) our robustness to variation in these rates across the genome. We also examined values of Garud et al.’s (2015) *H*_12_ and *H*_2_/*H*_1_ within windows classified by S/HIC as hard, soft, or neutral, noting that as expected, *H*_12_ is higher in sweeps than neutral regions, while *H*_2_/*H*_1_ is higher for soft sweeps than hard sweeps (supplementary fig. S2). Below, we begin with a brief overview of the broad patterns of adaptation we observe across populations, before discussing genomic features and biological pathways with a strong enrichment of selective sweeps, as well as compelling novel candidates for recently completed selective sweeps.

### The majority of sweeps in humans resemble selection on standing variation

We found a total of 1,927 distinct selective sweeps merged across all six populations (Methods). 190 (9.9%) of these are present in all populations, 59 (3.1%) are shared among the African populations, 71 (3.7%) are shared among the non-African populations, and 701 (36.4%) are population-specific (supplementary table S1). The remaining 906 (47.0%) sweeps were present in more than one population but do not fit into any of the categories above. We observe that across populations, the vast majority (1,776, or 92.2%) of sweeps were classified as soft, and note that this trend does not change qualitatively as we impose increasingly strict posterior probability thresholds before assigning a class label to a given window (supplementary table S2; Methods). These events may represent soft sweeps on standing genetic variants that our classifier was trained to detect, but we note that a similar signature can be created by a soft sweep resulting from recurrent origination of the adaptive allele(s), or by a *de novo* mutation that has been placed onto multiple haplotypes by allelic gene conversion events (see Discussion).

Although hard sweeps appear to be quite rare globally, the fraction of hard sweeps is significantly higher in non-African than African populations (table 1). For example, when comparing PEL to GWD, we observe a significantly higher fraction of hard sweeps in PEL (4.7% versus 1.6%; *p*=0.05). For each other African vs. non-African comparison we see an even greater (and more significant) disparity. Further, we observe a suggestive correlation between the fraction of sweeps in a population that were classified as soft and the harmonic mean of its population size within the last 4*N* generations (Pearson’s *ρ*=-0.96; Methods). Though taken at face value this correlation appears to be highly significant, we note that due to the six populations’ shared evolutionary history a statistical test of this correlation would be invalid.

**Table 1:**
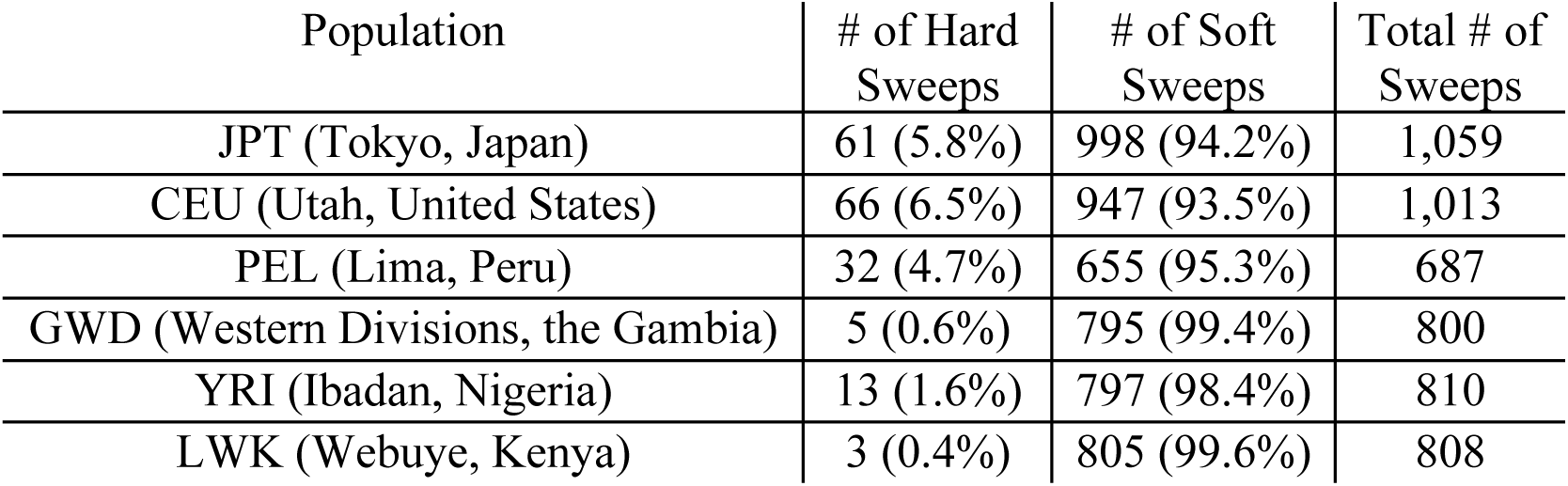
Number of sweeps of each type detected in each population sample.

| Population | # of Hard Sweeps | # of Soft Sweeps | Total # of Sweeps |
| --- | --- | --- | --- |
| JPT (Tokyo, Japan) | 61 (5.8%) | 998 (94.2%) | 1,059 |
| CEU (Utah, United States) | 66 (6.5%) | 947 (93.5%) | 1,013 |
| PEL (Lima, Peru) | 32 (4.7%) | 655 (95.3%) | 687 |
| GWD (Western Divisions, the Gambia) | 5 (0.6%) | 795 (99.4%) | 800 |
| YRI (Ibadan, Nigeria) | 13 (1.6%) | 797 (98.4%) | 810 |
| LWK (Webuye, Kenya) | 3 (0.4%) | 805 (99.6%) | 808 |

Comparing our results to those of previous scans we find that 519 of S/HIC’s sweep calls (26.9%) have previously been identified according to dbPSHP, a database of candidate regions for recent positive selection across human populations (Li et al. 2013). This accounts for 10.9% of the loci in the dbPSHP set (ignoring regions not classified by S/HIC). The remaining 1,408 sweeps called by S/HIC (73.1% of calls) represent potentially novel selective sweeps. There are several possible explanations for the modest overlap between our set of sweep candidates and those in dbPSHP. First, the sweep candidates in dbPSHP have been identified by a variety of methods, some of which are designed to detect selective scenarios other than completed sweeps (e.g. partial sweeps, spatially varying selection). Second, when comparing results from methods designed to detect the same type of sweeps, the intersection between studies is often fairly small (Akey 2009). Although most scans undoubtedly recover a large number of true selective sweeps, different methods may produce different false positives and false negatives, resulting in imperfect concordance between scans.

### Selective sweeps preferentially target genes involved in cancer and viral infection

Examining the locations of selective sweeps across the genome, we find that regions classified as selective sweeps are significantly overrepresented for both coding sequence and untranslated regions (*q*<0.05 in several populations for hard sweeps, and each population for putative soft sweeps; fig. 1A, B; supplementary table S3), relative to data sets with permuted classifications (see Methods). Enrichment for transcription factor binding sites was less pronounced, and only significant in soft sweeps for the three African populations along with PEL. The most striking result we observed was a dramatic enrichment of sweep windows for mutations in the COSMIC data set of somatic mutations that have been observed in cancer cells (Forbes et al. 2015) and may therefore play a role in tumor suppression/progression. Averaged across populations, the number of COSMIC mutations found in soft sweeps represents a 3.7-fold increase relative to that observed in permuted data sets; this enrichment was significant in each population, and peaked at 4.5-fold in PEL. For hard sweeps, this enrichment was 12-fold on average, reaching as high as 21-fold in CEU, though this was the only population for which the enrichment was statistically significant. We also observed a sizeable overrepresentation of genes encoding virus-interacting proteins (VIPs) curated by (Enard et al. 2016) in soft sweeps, with a 1.9-fold increase relative to permuted sets (averaged across populations). VIPs show a similar magnitude of enrichment in hard sweeps for some populations, but does not achieve significance at *q*<0.05.

**Figure 1:**
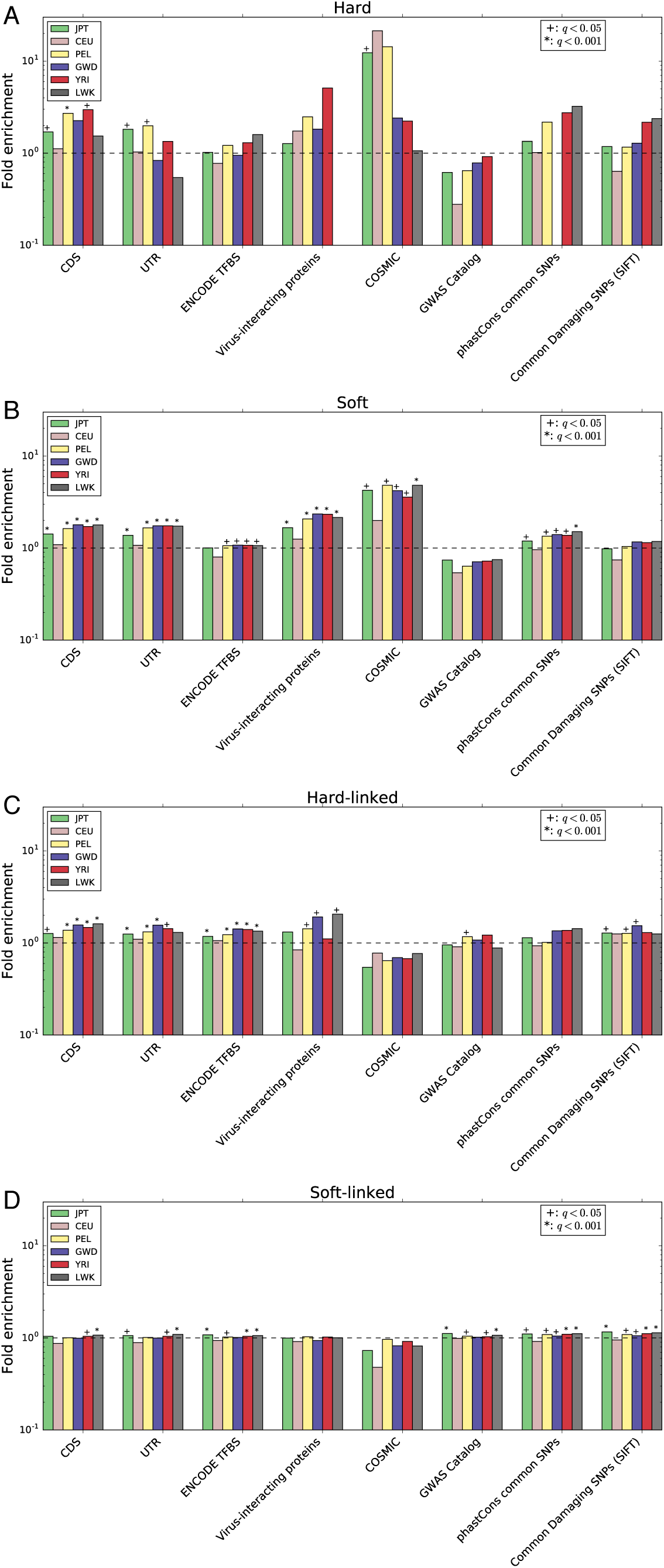
Enrichment of various annotation features in regions classified as sweeps or linked to sweeps relative. The fold enrichment is the ratio of the number of base pairs in the intersection between windows assigned to a given class and an annotation feature divided by the mean of this intersection across the permuted data sets (Methods). This was calculated separately for each population. (A) Enrichment of elements in windows classified as hard sweeps. (B) Same as A, but for soft sweeps. (C) Enrichment of elements in windows classified as affected by linked hard sweeps (D) Linked soft sweeps.

### Selective sweeps increase linked deleterious variation

Because S/HIC not only detects selective sweeps, but also attempts to identify regions of the genome that appear to be linked to recent sweeps, our classifications allow us to examine the effect of linked selection in a principled way. We found that while a minority of genomic windows were classified as selective sweeps (7.6% on average across all populations), a large fraction of windows were classified as linked to a completed selected sweep, either hard or soft (56.4% on average). These estimates range from 41.5% in JPT to 74.0% in GWD (fig. 2).

**Figure 2:**
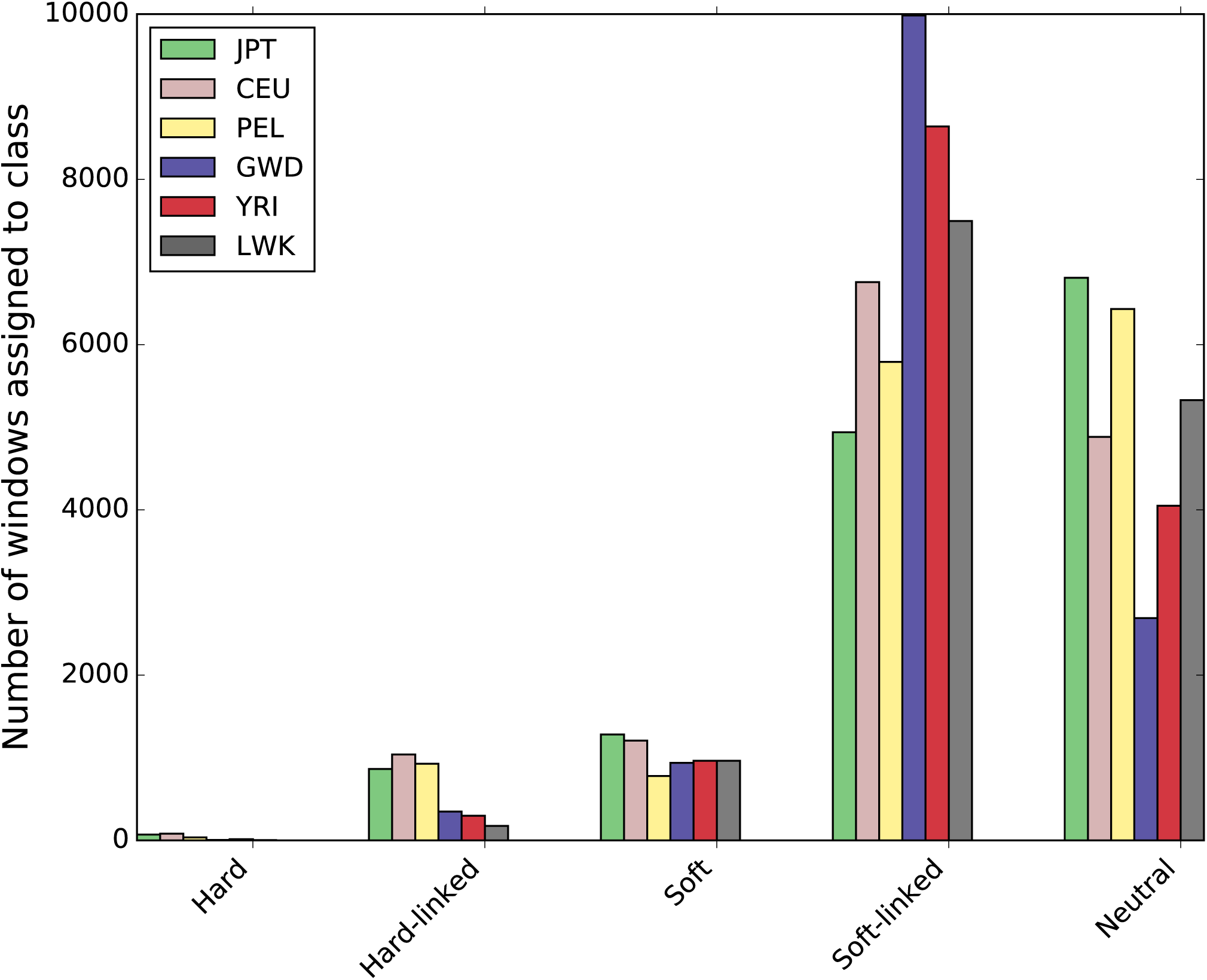
The number of windows assigned to each class by S/HIC in each population.

We also asked whether selective sweeps have a detectable impact on linked deleterious variation. As beneficial alleles increase in frequency in a population, they may carry along with them linked deleterious polymorphisms as hitchhikers, potentially increasing the frequency of deleterious variants over what would be expected given mutation-selection-drift equilibrium (Birky and Walsh 1988; Hartfield and Otto 2011). To this end we asked whether relatively common candidate deleterious mutations were enriched in regions classified as either hard-linked or soft-linked. Indeed, we observed a fairly subtle but significant overrepresentation of SNPs with derived allele frequencies of at least 0.01 but predicted to be damaging by SIFT (Kumar et al. 2009) in both the hard-linked (mean enrichment across populations: 1.3-fold) and soft-linked (mean enrichment: 1.1-fold) classes for most populations (fig. 1C, D; supplementary table S3). We find a similar enrichment in these sweep-linked classes of common SNPs in regions inferred to be conserved across primates according to phastCons (Siepel et al. 2005). Phenotype-associated variants from the GWAS catalogue (Welter et al. 2014) were also significantly overrepresented sweep-linked regions in several populations (Fig 1C, D).

### Sexual reproduction, the central nervous system, and immunity are targets of recent sweeps

In order to determine if positive selection preferentially acts on particular organismal functions, we asked which Gene Ontology (GO) terms were enriched in our sweep calls relative to the permuted data (Methods). In soft sweeps, we found a sizeable and significant enrichment (*q*<0.05) of terms related to sperm development, structure, and function. For example, “spermatogenesis” (4.4-fold enrichment averaged across populations), and “sperm-egg recognition” (3.9-fold enrichment on average) were enriched in soft sweeps in several populations. We also observed an overrepresentation of genes involved in the “glutamate receptor signaling pathway” in our soft sweep sets for each population (4-fold mean enrichment). Glutamate receptors are the primary excitatory neurotransmitter in the central nervous system, and important for both proper brain development and function (Luján et al. 2005). Indeed, soft sweeps are enriched for “central nervous system development” in multiple populations (1.6-fold mean enrichment). Numerous GO terms related to immune response, especially adaptive immunity, as well as KEGG pathways related to immunity and cancer progression/tumor suppression were also significantly enriched among soft sweeps (see supplementary table S4 for full list).

### Positive selection on interacting gene pairs

We examined three types of gene interaction networks: protein-protein interactions (PPIs), transcription factor-target gene interactions, and genetic interactions where one gene modifies the effect of another (Methods). Interestingly, we observed a dramatic enrichment of sweeps in genes that encode proteins that physically interact with one another (fig. 3A–B): if a gene overlapped a window classified as a soft sweep, genes that interact with this gene were on average 3.3 times more likely to overlap a putative soft sweep than expected by chance (*p*<0.0001 for each population; fig. 3B). Despite the smaller number of candidate regions, we found a significant enrichment for PPIs in hard sweeps, though this was only significant in for non-African populations (4.0-fold enrichment averaged across populations; *p*<0.05 in CEU, JPT, YRI; fig. 3A). For transcription factor-target interactions, we observe no overrepresentation of soft sweeps, but a significant enrichment of hard sweeps in non-African populations (*p*<0.05 for each; 8.5-fold enrichment on average; fig. 3C–D). There were no populations exhibiting an overrepresentation of pairs of genes with genetic interactions and experiencing sweeps of either type (fig. 3E–F).

**Figure 3:**
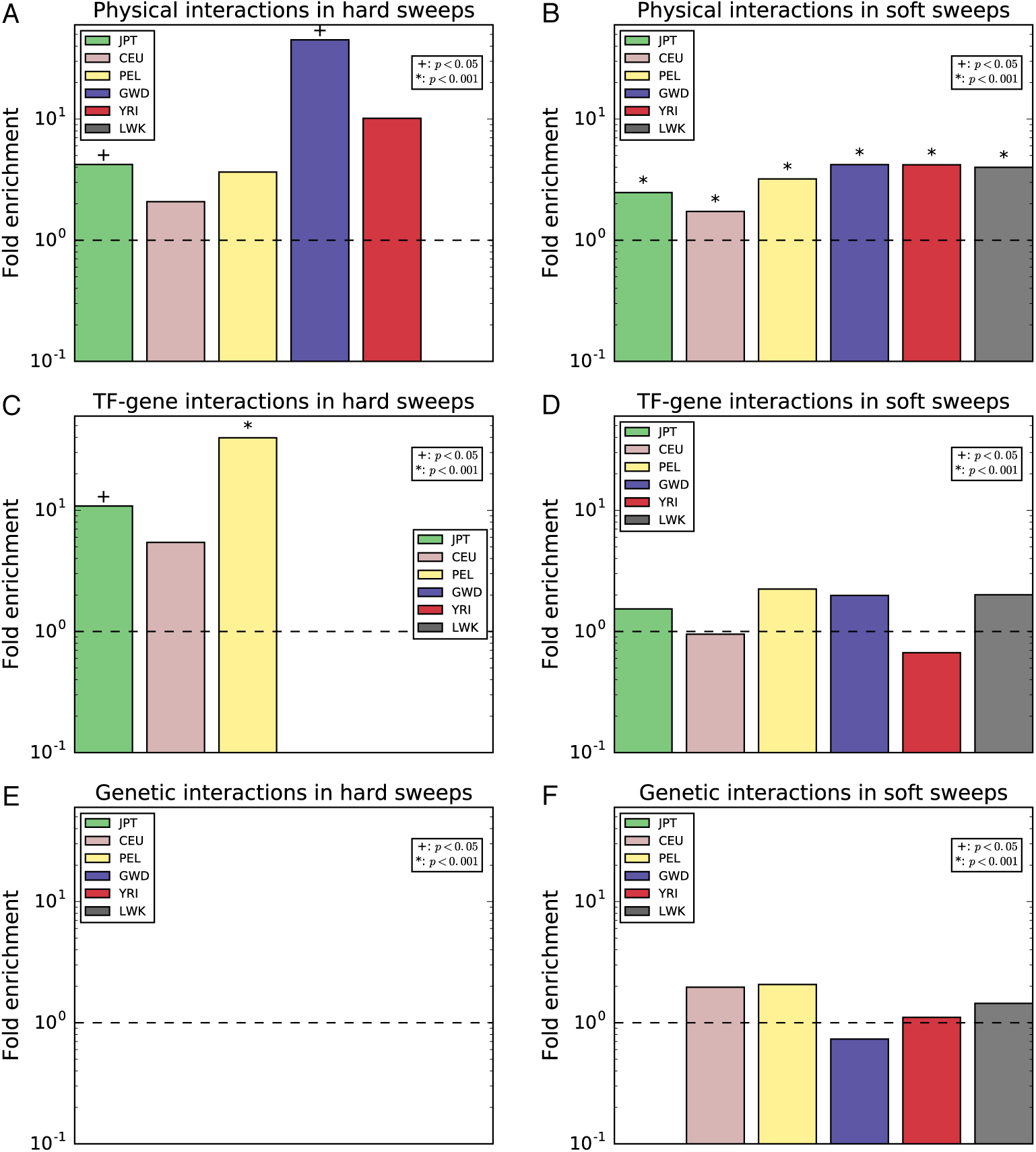
Enrichment of pairs of interacting genes each falling within a window classified as a sweep. The fold enrichment is the ratio of the number of pairs of interacting genes overlapping a window classified as a sweep of a given type divided by the mean of this number across the permuted data sets (Methods). This was calculated separately for each population. When no pairs of interacting sweep genes were observed in our true data set or a population, no bar was drawn. (A) Enrichment of pairs of genes encoding protein products that physically interact with each other (data from BioGRID) and both overlap hard sweep windows. (B) Same as A, but for soft sweeps. (C) Enrichment of pairs of genes, one of which is encodes a transcription factor that affects expression of the other (data from ORegAnno), where both overlap hard sweep windows. (D) Same as D, but for soft sweeps. (E) Enrichment of pairs of genes for which a genetic interaction has been observed (data from BioGRID) and both overlap hard sweep windows. (F) Same as E, but for soft sweeps.

### Examples of novel selective sweep candidates

In this section we describe several sweep candidates that exemplify the set of sweeps, and functions of putative targets of selection, that we were able to detect. As discussed above, our sets of sweeps were highly enriched for glutamate receptor-encoding genes. In supplementary fig. S3, we show a sweep candidate region on chromosome 4 that encompasses the glutamate receptor gene *GRIA2*. This sweep was previously detected in non-African populations by Pickrell et al. (2009), who did not find any evidence of selection in Africa. However, S/HIC infers that this region has experienced a soft sweep that is found in GWD and YRI, as well as the non-African populations. Consistent with this, Europeans, Asians, and African populations show a reduction in *π*, a trough in Tajima’s *D* (Tajima 1989), and a peak in Nielsen et al.’s SweepFinder composite likelihood ratio (CLR) test statistic, which captures regions that appear to be at the epicenter of the spatial skew in the SFS expected around sweeps (Nielsen et al. 2005b). Intriguingly, *GRIA2* interacts with the *GRID2* glutamate receptor gene (Kohda et al. 2003), which itself is classified as a soft sweep in CEU, LWK, PEL, and GWD. The remaining glutamate receptors overlapping identified sweeps are *GRIA4*, *GRID1*, *GRIK1*, *GRIK3*, *GRM2*, and *GRM7*. Of these genes, *GRIA4* and *GRID2* were shown by Liu et al. (2012) to have evolved a human-specific developmental expression profile.

Fig. 4 shows a region on chromosome 9 that exhibits strong evidence of a previously undetected hard sweep in each of our six populations. This region contains several members of the spermatogenesis associated 31 gene family: *SPATA31B1*, *SPATA31D1*, *SPATA31D3*, and *SPATA31D4*. Across populations this region shows dramatic valleys in *π* and Tajima’s *D*, as well as an elevated CLR near the center of the sweep window. These genes are highly testis-specific according to data from the GTEx project (Lonsdale et al. 2013), and male mice are infertile when lacking Spata31, another member of these gene family (Wu et al. 2015). fig. 4 also shows that each of these genes overlaps a cluster of non-repetitive piRNAs (data from piRBase; Zhang et al. 2014). Also near this region is *DDX10P2*, which GENCODE annotates as a processed pseudogene (Pei et al. 2012). *DDX10P2*, which is located at the center of the CLR peak for CEU, is expressed with a high degree of testis-specificity according to GTEx data, similar to the neighboring *SPATA31* genes. A BLAT search (Kent 2002) revealed that this putative pseudogene exhibits 99.5% sequence identity to the orthologous sequence in chimpanzees. The parent gene of *DDX10P2*, *DDX10*, is expressed in many tissues, but shows highest expression in the testis.

**Figure 4:**
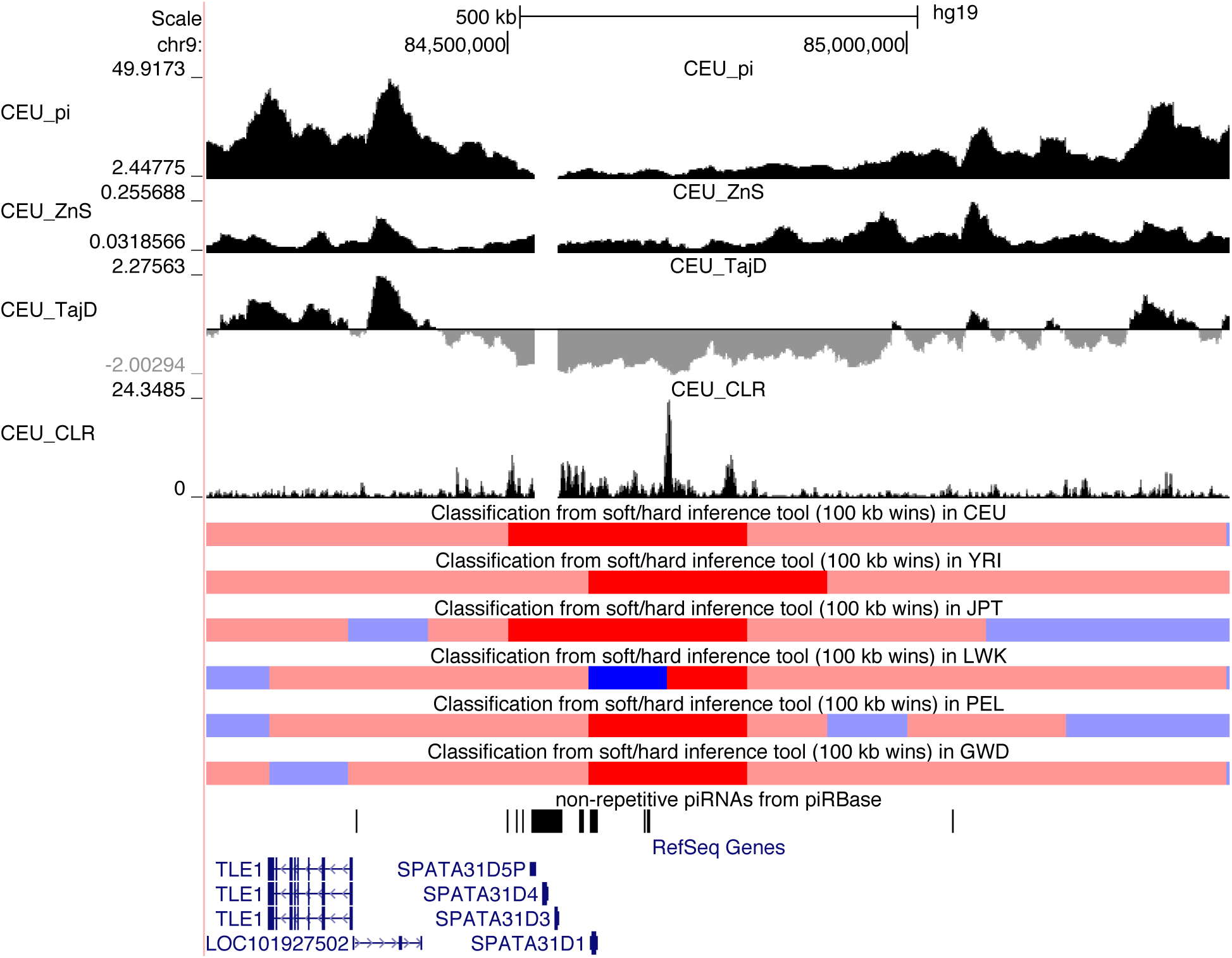
Hard selective sweep near several *SPATA31* spermatogenesis-associated genes. The S/HIC classification tracks show the raw classifier output for each population (red=hard sweep, blue=soft sweep, light red=hard-linked, light blue=soft-linked, black=neutral). We also show the values of various population genetic summary and test statistics (*π*, Tajima’s *D*, Kelly’s *Z*_*nS*_, and the SweepFinder composite likelihood ratio, or CLR). To avoid clutter, we only show statistics from CEU.

On chromosome 11 we detected what appear to be several novel soft sweeps present in and upstream of *CADM1* (cell adhesion molecule 1; fig. 5), one of which is present in each population. This gene is essential for spermatogenesis in mice (Van Der Weyden et al. 2006), and is also a tumor suppressor that is hypermethelated in various cancers (Kuramochi et al. 2001; Allinen et al. 2002; Fukuhara et al. 2002), as it works with the adaptive immune system to suppress metastasis (Faraji et al. 2012). *CADM1* is also active in the brain where it is involved in synaptic adhesion and has been linked to autism (Zhiling et al. 2008; Fujita et al. 2010). *CADM1* forms a complex with two other genes: the GABA receptor *GABBR2*, which has a soft sweep in YRI, and *MUPP1*, which has a soft sweep found in each population; this complex appears to localize to Purkinje cell dendrites (Fujita et al. 2012). Thus, this example encompasses many of the functions that we find are highly enriched across our sweep sets: adaptation in multiple interacting genes (one of which is a neurotransmitter), spermatogenesis, and tumor suppression (via adaptive immunity).

**Figure 5:**
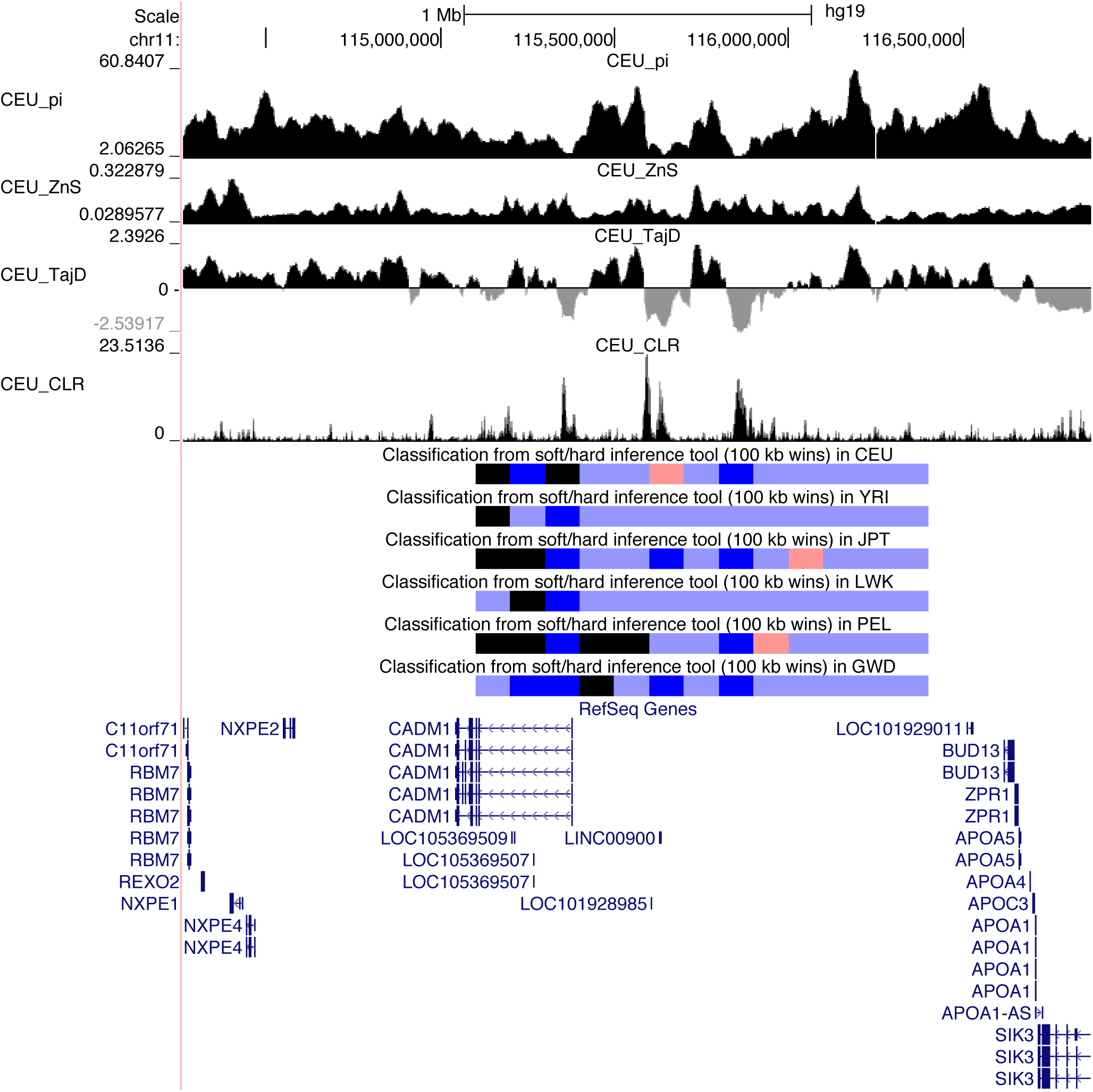
Soft selective sweeps near *CADM1*. The same tracks are shown as in Figure 4.

## DISCUSSION

Understanding the history of human adaptation at the genetic level is a central goal of population genomics and human evolutionary biology. Accordingly, since the completion of the human genome assembly (Lander et al. 2001) and subsequent proliferation of population genomic data, numerous genome-wide scans for selection have been conducted using differing methodologies (Sabeti et al. 2002; Voight et al. 2006; Sabeti et al. 2007; Pickrell et al. 2009; Field et al. 2016). The majority of these studies searched primarily for partial selective sweeps—the signature of a beneficial mutation currently sweeping through a population (see Williamson et al. 2007 for a notable exception)—and rightly so, as these sweeps can reveal the targets of ongoing adaptation in human populations. However, because the sojourn of an adaptive mutation to fixation should be rapid (e.g. on the order of 400 generations, assuming *N*=10^4^ and a moderately strong selection coefficient of *s* = 0.05, and 4000 generations for *s*=0.005), the success of efforts to detect ongoing selection implies the presence of a larger number of recently completed sweeps. We have therefore focused on completed sweeps in order to complement previous studies and to construct a more comprehensive catalogue of the loci underpinning recent human adaptation. Using a powerful and robust machine learning method that we have recently introduced (S/HIC; Schrider and Kern 2016) for finding completed selective sweeps, we performed a genome-wide search for the targets of recent positive selection in six human populations. Furthermore, we sought to determine the mode of positive selection, distinguishing between selection on *de novo* mutations and on previously standing variation.

### Soft sweeps dominate human adaptation

Perhaps our most consequential result is the finding that the majority of our candidate sweeps resemble soft sweeps on standing variation. This result implies that adaptation in humans may not be mutation-limited (Gillespie 1991; Karasov et al. 2010): rather than waiting for a novel mutation to arise, human populations may often be able to respond via selection on previously segregating polymorphisms, thereby more rapidly responding to novel environmental challenges. This may be surprising given the apparently small effective population size and low nucleotide diversity levels in humans. However, if the mutational target for the trait to be selected on is fairly large, then the probability of a population harboring a mutation affecting that trait may be appreciable.

While soft sweeps appear to be the dominant mode of selection globally, there is a significant increase in the proportion of putative hard sweeps in non-African populations relative to African populations. This is consistent with theoretical expectations, as larger populations have more standing variation for selection to act on (Hermisson and Pennings 2005). Moreover, the human migration out of Africa was associated with a severe population bottleneck (Marth et al. 2004; Fagundes et al. 2007). Soft selective sweeps may be “hardened” by a reduction in population size, which can result in the stochastic loss of some genetic backgrounds harboring the adaptive allele so that only a single haplotype reaches fixation (Wilson et al. 2014). Thus, though one might expect selection on segregating neutral or nearly neutral variation when a population enters a new environment with novel selective pressures, if the migration event is accompanied by a bottleneck then the population may experience a somewhat counterintuitive increase in the proportion of hard sweeps. Moreover, the causal relationship between population size and mode of adaptation may not be unidirectional. As Orr and Unckless (2014) have shown in the context of evolutionary rescue, when faced with a changing environment, a population which does not harbor standing variation that is beneficial may experience a more protracted decline in size while it waits for an adaptive *de novo* mutation.

Our genome-wide results amplify results of earlier studies that by design have tried to infer the mode of adaptation in a smaller number of targeted loci. For instance Peter *et al.* (Peter et al. 2012) attempted to infer the mode of adaptation among 7 loci previously identified to be under selection in human populations. They report that half of the loci that they could confidently classify supported selection on standing variation. In *Drosophila melanogaster,* when looking among strong outliers of haplotype homozgosity, Garud *et al.* (2015) found that patterns of variation in those regions were consistent with recent soft selective sweeps. Our finding, that the vast majority of sweeps in human populations are soft sweeps, thus underscores the ubiquity of selection from standing variation in natural populations. Indeed it seems plausible that adaptation from standing variation might be the rule, rather than the exception.

There are two caveats affecting our ability to discriminate between selection on standing variation and on *de novo* mutations. First, while we have trained our classifier to detect soft sweeps on previously segregating mutations, soft sweeps may also occur via recurrent mutation to the adaptive allele (Pennings and Hermisson 2006b, a). Though there are some qualitative differences between these two models of soft sweeps (Berg and Coop 2015; Schrider et al. 2015), these are fairly subtle in comparison to the differences between the other models we consider. Thus, our classifiers may have sensitivity to both types of sweeps. If this is so, then some of the soft sweeps that we detect may result from recurrent mutation. Additionally, gene conversion during a sweep can transfer the adaptive mutation on to new genetic backgrounds (Jones and Wakeley 2008), thereby “softening” the sweep (Schrider et al. 2015). This implies that selection on a single *de novo* mutation could sometimes appear to be a soft sweep in our classification. In any case, our finding that most sweeps in humans do not appear to be hard sweeps underscores the importance of using methods that are sensitive to soft sweeps.

### Extensive impact of linked positive selection

Our analysis demonstrates that the impact of linked positive selection on genetic variation is considerable, with roughly half of the genome classified by S/HIC as being influenced by a nearby sweep. This result has important implications for efforts to infer demographic histories from patterns of genetic polymorphism, as most inference methods hinge on the assumption of neutrality. Indeed, we have recently shown that linked positive selection has the potential to severely confound demographic inferences (Schrider et al. 2016). Similarly, Ewing and Jensen (2016) have found that background selection (Charlesworth et al. 1993) can also bias demographic estimates. One strategy is to use only those polymorphisms that are distant from genes and conserved noncoding elements to mitigate these effects (Gazave et al. 2014). One could further supplement such an approach by using S/HIC to directly ask which intergenic regions are unaffected by hitchhiking in order to further diminish the bias introduced by linked selection. We note that the putatively neutrally evolving regions found in this study can be obtained from our raw classification output (available at https://github.com/kern-lab/shIC/tree/master/humanScanResults).

If linked positive selection affects much of the genome, then that implies that the frequencies of many neutral or weakly deleterious mutations may be altered by genetic draft (Gillespie 2000). That is to say, deleterious mutations that happen to reside on chromosomes that begin to sweep may be able to reach higher frequencies than expected from mutation-selection-drift equilibrium. Consistent with this, we observe a slight but significant excess of potentially deleterious polymorphisms in windows classified as linked to selective sweeps. Previously, Chun and Fay (2011) found evidence that the ratio of deleterious to neutral polymorphisms is elevated in sweep regions, concluding that hitchhiking carries linked deleterious variants to higher frequencies. Our finding that SNPs from the GWAS catalogue are also enriched regions linked to selective sweeps lends further support to this hypothesis. Indeed, several compelling examples of hitchhiking mutations known or suspected of causing disease have been described in the literature (Helgason et al. 2007; Chun and Fay 2011; Huff et al. 2012). Moreover it seems that the phenomenon of deleterious alleles hitchhiking along with strongly beneficial alleles is not restricted to humans: a recent study also uncovered evidence that selection during domestication increased the frequency of deleterious polymorphisms in dogs (Marsden et al. 2016).

### Targets of recent human selective sweeps

Our catalogue of sweep candidates allowed us to characterize the biological functions that are overrepresented in sweeps. Notably, we found a strong excess of spermatogenesis genes within sweep regions, a phenomenon previously observed by Voight et al. (2006). This signature may be a result of sexual selection, sexual conflict, and/or sperm competition (Swanson and Vacquier 2002). We also observed a significant enrichment of cancer-related genes among our sweep candidates. Nielsen et al. (2005a) found a similar enrichment of candidate genes under selection related to cancer when examining protein divergence between humans and chimpanzees. These authors found that some of these genes are also involved in spermatogenesis (much like our *CADM1* example), and concluded that genomic conflict between tumor suppression and the advantage of avoiding apoptosis during spermatogenesis may explain the selection on cancer genes. An alternative (and non-mutually exclusive) explanation is that the increase in longevity along the human lineage has created an immense selective pressure to reduce the rate of cancer progression by orders of magnitude (Nunney and Muir 2015).

We also observed a significant excess of glutamate receptor genes targeted by sweeps, suggesting that these loci may underlie some of the dramatic neurological changes that have occurred along the human lineage. Consistent with this, we previously found evidence suggesting some of these glutamate receptor genes (along with other neurotransmitters) may have recently gained novel regulatory elements in humans (Schrider and Kern 2015; Meyer et al. 2017). The most striking examples of glutamate receptors experiencing sweeps are *GRIA2* and *GRID2*, which show strong signatures of selection in multiple populations and physically interact with one another. The action of positive selection on multiple members of the protein complex appears to be a general phenomenon (fig. 3). For a more in-depth examination of positive selection in the PPI network, see Qian et al. (2015), who found that genes in candidate regions for positive selection were more likely to lie close together in the PPI network.

## Conclusions

Our investigation has revealed several valuable insights into the adaptive process in human populations. The success of our approach exemplifies the potential of machine learning methods to elucidate the adaptive process in humans and other species (Fan et al. 2016). To date several machine learning methods have been devised to detect selective sweeps (Pavlidis et al. 2010; Lin et al. 2011; Ronen et al. 2013; Pybus et al. 2015; Sheehan and Song 2016), and they tend to substantially outperform more traditional approaches (see Schrider and Kern 2016). We suspect that machine learning could be used to make important inroads in answering a variety of evolutionary questions.

Finally, Hernandez et al. (2011) argued that hard selective sweeps might be rare in human populations, and instead suggested that the majority of adaptation might be a consequence of selection on standing variation or selection on polygenic traits. We here find direct evidence that indeed this is the case—the vast bulk of human adaptation is occurring as a consequence of soft sweeps. Our observation thus reconciles Hernandez et al.’s findings with those of Enard et al., who conclude that the reduction in diversity around amino acid substitutions is caused by widespread selective sweeps (Enard et al. 2014). Moreover, while our scan leveraged a method that performs very well in detecting both hard and soft sweeps, it was not trained to detect cases of polygenic selection (e.g. Berg and Coop 2014). It is fair to assume that a large majority of phenotypes are determined by multiple loci, thus polygenic selection should be expected to be common. If that were the case, then it could very well be that an even larger portion of genetic variation is influenced by natural selection and its linked effects throughout the genome.

## METHODS

### Sequence and annotation data

We downloaded phased genotype data from Phase 3 of the 1000 Genomes Project (Auton et al. 2015). This data set consists of 26 population samples from Africa, East Asia, South Asia, Europe, and the Americas. We wished to include only populations where the influence of admixture/migration on genetic variation appeared to be minimal, while still allowing us to characterize selection across multiple continents. We therefore chose to scan the following populations for selective sweeps: the GWD (Gambians in Western Divisions in The Gambia) and YRI (Yoruba in Ibadan, Nigeria) populations from West Africa, LWK (Luhya in Webuye, Kenya) from East Africa, JPT (Japanese in Tokyo, Japan) from Asia, CEU (Utah residents with Northern and Western European Ancestry) from Europe, and PEL (Peruvians from Lima, Peru) from the Americas. Examining Auton et al.’s results from running ADMIXTURE (Alexander et al. 2009), we see that for most values of *K*, each of these populations appears to correspond primarily to a single ancestral population rather than displaying multiple clusters of ancestry (see Extended Data Figure 5 from Auton et al. 2015). One exception may be the PEL population, but among the highly admixed American samples it appears to exhibit the smallest amount of possible mixed ancestry (for most values of *K*), so we retained this population in order to have some representation from the Americas. We opted not to examine any South Asian population, as for each of these samples ADMIXTURE inferred evidence of ancestry from three or more ancestral populations.

We downloaded numerous annotation data sets containing genomic features to test for enrichment/depletion of selective sweeps and perform other downstream analyses. These included GENCODE gene model release 19 (Harrow et al. 2012) including pseudogenes (Pei et al. 2012), virus-interacting proteins from Enard et al. (2016), enhancers gained or along the human lineage since diverging from Old World monkeys (Cotney et al. 2013), and SIFT’s (Kumar et al. 2009) predictions of damaging amino acid polymorphisms from dbNSFP version 3.2a (Liu et al. 2016). We obtained Gene Ontology (GO) annotations from ENSEMBL release 75 (Yates et al. 2016). We also downloaded coordinates of previously identified selective sweeps from dbPSHP (Li et al. 2013).

We used the UCSC Table Browser (Karolchik et al. 2004) to obtain the following data sets: phenotype-associated SNPs from the GWAS Catalog (accessed Apr 12, 2016; Welter et al. 2014), ClinVar pathogenic SNPs and indels ≤ 20 bp in length (Apr 26, 2016; Landrum et al. 2016), COSMIC somatic mutations in cancer (accessed Feb 25, 2014; Forbes et al. 2015), phastCons elements conserved across primates (accessed Jun 2, 2013; Siepel et al. 2005), ENCODE transcription factor binding sites version 3 (accessed Aug 25, 2013; Dunham et al. 2012), tables of genes and SNPs implicated in Mendelian phenotypes from OMIM (accessed May 2, 2016; Amberger et al. 2015), and KEGG pathway annotations (accessed Apr 27, 2016; Kanehisa et al. 2015). For each of these data sets we used GRCh37/hg19 coordinates.

In order to examine the prevalence of selective sweeps within interacting gene networks, we downloaded physical and genetic interactions from BioGRID version 3.4.136 (Chatr-Aryamontri et al. 2015). Our set of genetic interactions consisted of those annotated as “synthetic genetic interaction defined by inequality,” “suppressive genetic interaction defined by inequality,” or “additive genetic interaction defined by inequality.” Physical interactions included those annotated as “direct interaction,” “association,” or “physical association.” We extracted transcription factor-target interactions from ORegAnno (accessed Dec 22, 2015; Griffith et al. 2008), retaining only interacting pairs where the ENSEMBL gene identifier were provided for both genes in order to avoid ambiguity.

### Building classifiers to detect selective sweeps

To detect sweeps we used S/HIC (https://github.com/kern-lab/shIC), a machine learning approach we previously described and showed to be remarkably powerful and robust to non-equilibrium demography (Schrider and Kern 2016). Briefly, the S/HIC machine learning approach leverages spatial patterns (along a genome) of a variety of population genetic summary statistics to classify genomic windows as being the target of a completed hard sweep (hard), being closely linked to a hard sweep (hard-linked), a completed soft sweep (soft), linked to a soft sweep (soft-linked), or evolving neutrally (neutral). While this classification approach allows inference when considering a large number of features jointly, it necessitates training from a large number of data instances known to belong to each class. Because the number of genomic windows known to belong to each our five classes is limited, we must rely on simulation to generate our training data. To this end we used the program discoal (Kern and Schrider 2016) to simulate large chromosomal regions, subdivided into 11 sub-windows. Training examples for the hard class experienced a hard sweep in the center of the central sub-window (i.e. the 6^th^ window), while examples for the hard-linked class experienced a hard sweep in the center of one of the remaining sub-windows (selected randomly). Analogous simulations with soft sweeps were generated for the soft and soft-linked classes, respectively. Finally, neutrally evolving examples did not experience any selective sweep.

We sought to train a classifier for each population under a demographic model that offers a better approximation to the population size history than the standard neutral model. For this we used Auton et al.’s (2015) population histories inferred by PSMC (Li and Durbin 2011). The 1000 Genomes Project’s PSMC output did not contain estimates of *θ*, the population mutation rate parameter. Thus for each population we conducted a grid search by simulating genomic windows with the appropriate sample size under each demographic model with varying values of *θ*=4*NuL* (where *L* is the length of the locus, which we set to 100 kb); the grid of *θ* values raged from 10 to 250, examining multiples of 10. For each value of *θ*, we compared the values of *π* (Nei and Li 1979), 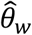 (Watterson 1975), 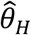 (Fay and Wu 2000), *H*_2_/*H*_1_ (Garud et al. 2015), and *Z*_*nS*_ (Kelly 1997) from 1000 simulations to those from 1000 randomly selected genomic loci (calculated as described below), calculating the mean of each statistic in the real and simulated datasets. We chose as the final values of *θ* that for which the sum of the percent deviations of the simulated from the observed means of each statistic was minimized. This estimate of *θ* allowed us to calculate estimated population sizes and times scaled by the number generations for each time point in the history inferred by PSMC. The harmonic mean of each population’s size was calculated by taking the estimated population size for each of the last 4*N* generations. We note that these models may not accurately capture the demographic histories of the populations we examined due to the confounding effects of positive (Schrider et al. 2016) and negative (Ewing and Jensen 2016) selection. However, because of S/HIC’s robustness to demographic misspecification, we do not expect this to severely impact our analysis (Schrider and Kern 2016).

For each population we simulated a total of 2000 regions for each of our five classes. For simulations involving sweeps, we drew the selection coefficient from *U*(0.005, 0.1), the sweep completion time from *U*(0, 2000), the initial selected frequency for soft sweeps from *U*(1/*N*, 0.2). We drew values of *θ* uniformly from a range spanning exactly one order of magnitude, specified so that the mean value of *θ* was equal to that estimated for the population as described above. We drew recombination rates from an exponential distribution with mean 1×10^−8^, truncated at triple the mean due to memory constraints. The simulation program discoal requires some of these parameters to be scaled by the present-day effective population size; we did this by taking the mean value of *θ* and dividing by 4*uL*, where u was set to 1.2×10^−8^ (Kong et al. 2012). The full command lines we used to generate 1.1 Mb regions (to be subdivided into 11 windows each 100 kb in length) for each population are shown in supplementary table S5. We also simulated 1000 test examples for each population in the same manner as for the training data.

In order to address the potential for purifying and background selection to confound our classifiers, we simulated additional test sets of 1000 genomic windows 1.1 Mb in length with varying arrangements of selected sites. In order to mimic patters of purifying/background selection expected in the human genome as closely as possible, for each of our 1000 replicates we randomly selected a 1.1 Mb window from the human genome and asked which sites were found within either a GENCODE exon (Harrow et al. 2012) or within a phastCons (Siepel et al. 2005) conserved element from the UCSC Genome Browser’s 100-way vertebrate alignment (Kent et al. 2002). Sites in the simulated chromosome corresponding to these functional elements in the human genome were labeled as “selected” in the simulations. In “selected” regions, 25% of all new mutations had no fitness effect, while the remaining 75% had a selection coefficient drawn from a gamma distribution with mean of −0.0294 and a shape parameter of 0.184 (the African model from Boyko et al. 2008). We limit fitness effects of new mutations to 75% in an effort to mimic coding regions of the genome. We note that this percentage may not be accurate for noncoding functional regions, though it is likely that some fraction of mutations in these regions is effectively neutral. All mutations outside of the selected regions were fitness-neutral. These simulations were performed for both our GWD and JPT demographic models using the fwdpy11 (https://github.com/molpopgen/fwdpy11) forward population genetic simulator (Thornton 2014), using the same mutation rates, recombination rates, and history of instantaneous population size changes as used in our coalescent simulations described in supplementary table S5. Feature vectors were then generated for each of these simulated test examples in the same manner as for our coalescent simulations. We also tested each population’s classifier against test sets generated by discoal with different fixed values of *θ* (but otherwise with the same parameterizations shown in supplementary table S5) in order to ensure that our approach was robust to uncertainty in the estimate of this parameter (supplementary fig. S4).

Our feature vector for each simulated region examined the spatial patters (following Schrider and Kern 2016) of each of the following statistics: *π* (Nei and Li 1979), 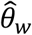 (Watterson 1975), 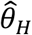 (Fay and Wu 2000), the number of distinct haplotypes, average haplotype homozygosity, Garud et al.’s (2015) *H*_12_ and *H*_2_/*H*_1_ statistics, *Z*_*nS*_, *ω* (Kim and Nielsen 2004), and the maximum frequency of derived mutations (Li 2011). Before calculating these summary statistics we masked a number of sites within each simulation by randomly selecting a 1.1 Mb region from our empirical windows sampled throughout the genome and masking the same regions in the simulated window as were masked in the genomic window (see below). Thus our simulated windows exhibit the same distribution of regions of missing data as the windows to which we applied our classifiers. We then used S/HIC to train extra-trees classifiers (Geurts et al. 2006), one for each population.

### Classifying genomic windows in each population

Having trained our classifiers, we then applied them to genomic data from the corresponding population. We inferred ancestral states of polymorphisms and masked inaccessible sites (whether polymorphic or not) in the same manner as described previously (Schrider and Kern 2016). We then used S/HIC to classify the central 100 kb sub-window of 1.1 Mb windows across the autosomes, while taking the stringent approach of omitting those for which any sub-window was less than 25% accessible, before sliding 100 kb downstream to examine the next window. We also removed windows where any of the three central sub-windows had a mean recombination rate of zero (using data from Kong et al. 2010). Importantly, for each retained 1.1 Mb window, we recorded the locations of all sites deemed inaccessible for use in masking our training data (see above). In total we classified 13,968 windows, accounting for 48.5% of the assembled autosomes. For our classifications we simply took the class that S/HIC’s classifier inferred to be the most likely one, but we also used S/HIC’s posterior class membership probability estimates in order to experiment with different confidence thresholds (results shown in supplementary table S2). For a given threshold, we required the sum of a windows’ hard and soft sweep posterior probabilities to be greater than or equal to the threshold before labeling the window as a sweep; the mode of the sweep was that corresponding to the greater posterior probability among the hard and soft sweep classes.

In order to count the number of distinct sweep candidates found within our set of populations, we simply merged all 100 kb windows classified as a sweep of either type that were located either at the exact same coordinates or adjacent to one another, repeating this until no more sweep regions could be merged. If all constituent windows were classified as soft, we counted the sweep as soft; otherwise we counted it as a hard sweep. We used a similar approach but examining classifications from only one population at a time in order to count the number of sweeps of each type in that population. If a gene found within a sweep window identified by S/HIC was not found in an entry of dbPSHP (Li et al. 2013), we referred to it as a novel sweep. Visualization of sweep candidates was performed using the UCSC Genome Browser (Kent et al. 2002), along with custom tracks showing values of various population genetic summary statistics and selection scan scores for the CEU, YRI, and JPT populations from the Human Positive Selection Browser (Pybus et al. 2013). Our classification results are available at https://github.com/kern-lab/shIC/tree/master/humanScanResults.

### Permutation tests for enrichment of annotation features in sweeps

To determine whether certain annotation features were enriched within any of our five classes, we carefully designed a permutation test to account for the subset of the genome that we examined with S/HIC, as well as the spatial correlation of S/HIC’s classifications (i.e. adjacent windows are especially likely to receive the same classification). Briefly, the permutation algorithm begins by examining our classification results for a given population and keeping track of the length of runs of consecutive windows assigned to each class. The permutation algorithm then selects a chromosome, and begins at its first classified window (i.e. not removed by data filtering). A run length and associated class assignment is then randomly drawn without replacement. This process continues until the end of the chromosome, and then another chromosome is selected until the end of the final chromosome is reached, at which point the permutation has been completed. We then repeated this permutation procedure 10,000 times for each population. Note that this process preserves the run length distribution of our classifications while permuting them across the set of genomic windows that had enough unmasked data to be included in our scan.

After constructing our permuted data sets, we conducted one-sided enrichment tests by counting the number of base pairs in the intersection between the S/HIC class of interest and the annotation feature of interest, and comparing this number to its distribution among the permuted data sets. The fraction of permuted data sets where this intersect was greater than or equal to that observed for the real data is the *p*-value. Because we tested each of S/HIC’s five classes for enrichment of a fairly large number of genomic features (supplementary table S3), we corrected for multiple testing using false discovery rate *q*-values following Storey (2002). When testing GO terms and KEGG pathways for enrichment, we considered only the hard and soft sweep classes, corrected for calculating *q*-values separately for each class.

We also asked whether the number of pairs of interacting genes both overlapping windows classified as sweeps was greater than in our permuted data sets. To ensure that our results were not inflated by the spatial clustering of interacting genes, we only counted interacting pairs overlapping sweep windows if they were separated by at least 10 Mb or on separate chromosomes. In addition, if we observed an interaction between two genes, *A* and *B*, that each overlapped sweeps, and a third sweep candidate gene, *C*, was found, to avoid redundancy we counted at most one interaction between *A* and *C* and *B* and *C*, even if *C* was found interact with both other genes. As with GO and KEGG terms, we only searched the hard and soft classes for enrichments before calculating one-sided q-values as described above.

## ACKNOWLEDGMENTS

We thank Matthew Hahn, David Enard, and three anonymous reviewers for feedback on the manuscript. D.R.S. was supported in part by NIH award no. K99HG008696. A.D.K. was supported in part by NIH award no. R01GM078204.

## SUPPLEMENTARY FIGURE AND TABLE LEGENDS

**supplementary fig. S1: Heatmaps showing the accuracy of our six classifiers on test data, one for each population.** On the *y*-axis, we show the location of the sweep relative to the classified window (i.e. the central sub-window), with the exception of the “Neutral” case where there is no sweep. The test data were simulated under the same demographic models used for training. On the *x*-axis we show the class inferred by S/HIC. A perfect classifier would infer “Hard” for 100% test instance where a hard sweep is in the focal sub-window (and analogously for soft sweeps), “Hard-linked” for 100% of cases where a hard sweep occurs elsewhere (and analogously for soft sweeps not located in the central sub-window), and “Neutral” for 100% of cases with no sweep. Both GWD and JPT also contain test results on a simulated set of examples of purifying/background selection. (A) Test results for CEU. (B) GWD. (C) JPT. (D) LWK. (E) PEL. (F) YRI.

**supplementary fig. S2: Histograms of *H*_12_ and *H*_2_/*H*_1_ within windows classified has hard sweeps, soft sweeps, or neutral for each population.**

**supplementary fig. S3: Soft selective sweep in *GRIA2*.** The same tracks are shown as in figs. 4 and 5.

**supplementary fig. S4: False positive and false negative rates on simulated test data with varying values of *θ*.** For each population, we used discoal to simulate 100 replicates for each combination of S/HIC’s five classes and three fixed values of *θ*. In these simulations all parameters other than *θ* had the same values as in supplementary table S5. “Medium” *θ* refers to the mean value of *θ* used for a given population’s training and testing simulations (Methods), while “Low” and “High” *θ* refer to one-half and double this value, respectively. Examples of the Hard-linked, Soft-linked, or Neutral classes that are classified as sweeps represent false positives, while Hard and Soft examples not classified as sweeps are false negatives.

**supplementary table S1: Number of sweeps found in each subset of populations.**

**supplementary table S2: Numbers of hard and soft sweeps found in each population when imposing various posterior probability thresholds to S/HIC’s classifications.**

**supplementary table S3: Enrichment of various sequence annotations in each S/HIC class.**

**supplementary table S4: Enrichment of annotation terms in hard and soft sweeps (only terms with *q*<0.05 for at least one sweep type in at least one population are shown).**

**supplementary table S5: Example command lines used to generate training data for each population, with a soft sweep occurring in the central sub-window.**

